# Local exclusion and regional decline of an endemic Galápagos tree species (*Psidium galapageium*) by an invasive relative (*P. guajava*)

**DOI:** 10.1101/2022.10.11.511772

**Authors:** Bryan Reatini, María de Lourdes Torres, Todd J. Vision

## Abstract

Invasive species can interact with native relatives in a variety of ways which may jeopardize their long-term coexistence. Here we show that interactions with an invasive species of guava (*Psidium guajava*) appear to be driving the local exclusion and regional decline of guayabillo (*Psidium galapageium*), a tree species endemic to the Galápagos archipelago. We find evidence consistent with recent historic exclusion of guayabillo from the highlands of San Cristóbal Island, signatures of ongoing demographic decline in sympatric populations at lower elevations, and evidence suggesting that the four coinhabited islands represent points along a time series of regional decline, with the extent of guayabillo decline depending on the date that guava was introduced to each island. Based on these results, we then use the percentage of guava cover surrounding guayabillo populations to target populations that are at imminent risk of exclusion to aid in prioritizing management targets.

## Introduction

Common guava (*Psidium guajava*) is among the most problematic and longest standing invasive species in the Galápagos islands (Laso et al., 2020; Rivas-Torres et al., 2018; Torres & Mena, 2018). Since its original introduction to San Cristóbal island in the late 1800s (Urquía et al., 2019), it has spread aggressively throughout the highlands of the four human inhabited islands (Laso et al., 2020; Rivas-Torres et al., 2018), where it has come into secondary contact with a related endemic, guayabillo (*P. galapageium*). Guayabillo is now absent from the highlands of San Cristóbal – where the guava invasion began and where guava still dominates today (Laso et al., 2020; Urquía et al., 2019) – despite guayabillo occurring in the highlands of all other islands within its archipelago-wide distribution (Figure S1,S3,S4). Guayabillo populations on San Cristóbal are also fragmented and have significantly less genetic variation than populations on the other inhabited islands (Urquía et al., 2020). Together, these observations suggest that guava may have excluded guayabillo from the highlands of San Cristóbal, and that the sympatric populations of guayabillo at lower elevations where guava has become established more recently may currently face a threat of extirpation.

Although many biotic and abiotic factors can contribute to demographic decline, there are reasons to suspect that interactions with guava might be driving guayabillo’s decline on San Cristóbal. First, land use patterns such as clear-cutting native forests for ranches and farmland are unlikely to be directly responsible for all of guayabillo’s decline since San Cristóbal has far less pasture and crop land relative to all other inhabited islands (Laso et al., 2020) – although this does not rule out an impact from the longer history of agricultural practices on this island. Second, guava is by far the most dominant invasive species by percentage of land cover in both the agricultural zone and surrounding Galápagos National Park – particularly where guayabillo is now absent (Laso et al., 2020; Rivas-Torres et al., 2018) (Figure 1B). Third, guava is reported to negatively impact a wide range of native species by way of competition and habitat alteration, including within other island ecosystems such as Hawaii (CABI, 2022).

**Figure 1.**
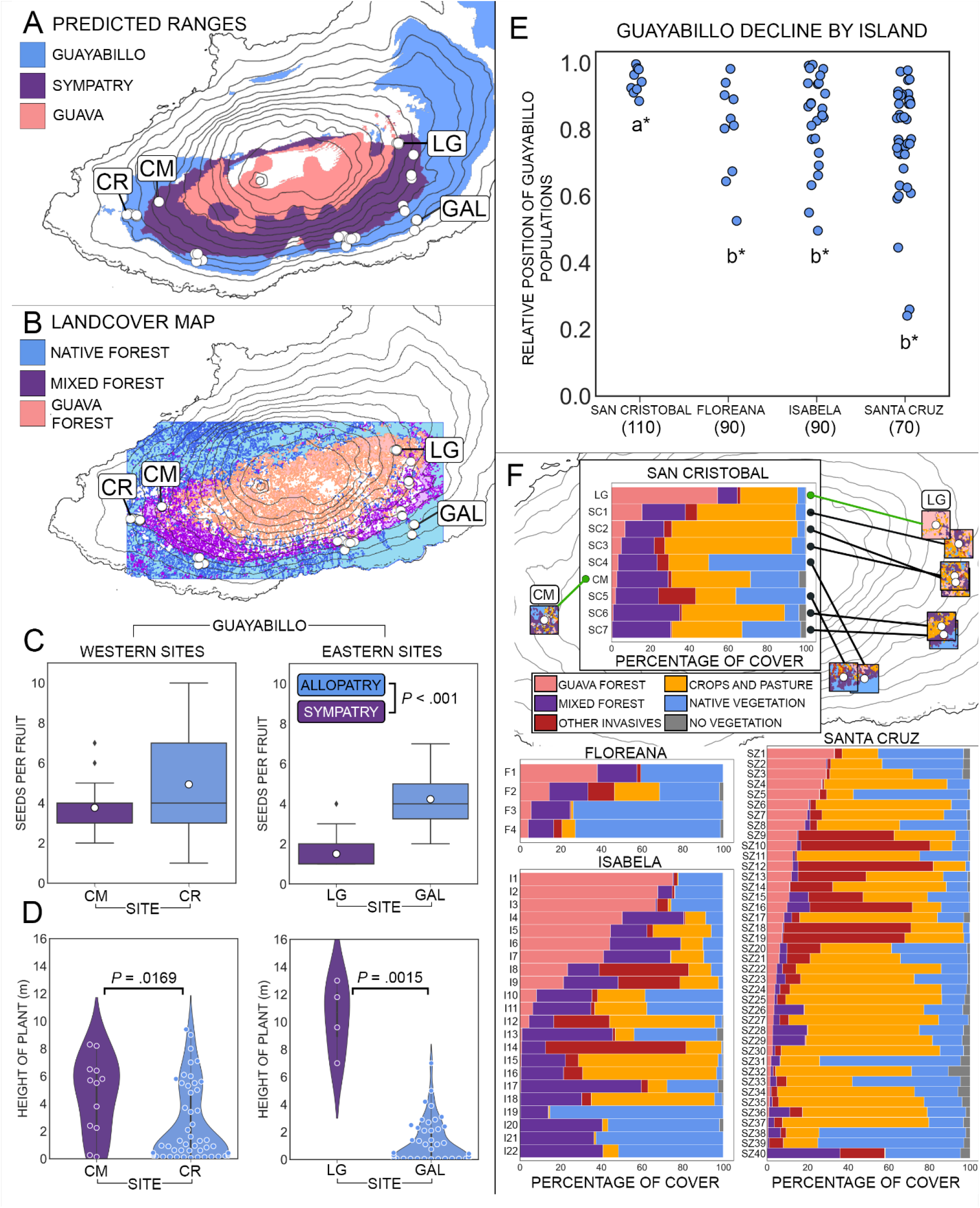
Evidence for guayabillo’s local exclusion on San Cristóbal and regional decline across the Galápagos archipelago. A) Species distribution models show that extant guayabillo populations on San Cristóbal (white points) occur along the edge of guava’s predicted range, despite much of the highlands being suitable for both species. B) Land cover maps from Laso et al. (2020) confirm that guava dominated forests cover much of the highlands of San Cristóbal, including a large proportion of predicted sympatry from A. Mixed forests contain a mix of native and introduced vegetation including both guava and guayabillo in some regions, whereas native forests contain predominantly native species including guayabillo in some regions. C) Guayabillo seed set is significantly lower in sympatry relative to allopatry for both western and eastern site pairs. Mean values of seed set are shown by white dots. Lower seed set in sympatric LG relative to sympatric CM is consistent with LG being in a region of higher guava density as seen in B and F. D) Using tree height as a rough proxy for age, guayabillo juveniles are rare in sympatry relative to allopatry for both western and eastern site pairs – consistent with reduced seed set in C. Demographic decline appears to be more advanced in sympatric LG than CM, which again is consistent with LG being in a region with higher guava density. E) The relative positions of guayabillo populations within predicted sympatry (blue points) reveal that exclusion is most advanced on San Cristóbal, least advanced on Santa Cruz, and intermediate for the two islands with intermediate dates of introduction. Letters signify significance groups. F) Percentages of land cover within 1 km^2^ quadrats surrounding guayabillo populations reveal considerable variation in guava cover (pink) and mixed forest cover (purple) throughout the archipelago, providing an opportunity to target the relatively few populations with very high guava cover for immediate conservation action. For San Cristóbal, populations are linked to their map positions, and the distribution of land cover is shown within each 1 km^2^ quadrat. Green lines indicate the sympatric study sites LG and CM to highlight the large difference in guava density between these the sites.

Here we test the hypothesis that invasive guava has excluded guayabillo from the highlands of San Cristóbal, and that guayabillo populations at lower elevations and on islands with more recent guava introductions are under threat of exclusion. Specifically, we first investigate whether the predicted and realized distributions of each species exhibit patterns consistent with a recent history of guayabillo decline due to guava encroachment. Next, we test whether remaining sympatric populations of guayabillo on San Cristóbal Island show evidence of demographic decline relative to allopatric populations. Given the results of these analyses, we then test whether islands with more recent guava invasions show weaker signs of regional decline for guayabillo populations. Finally, we rank sympatric guayabillo populations by the density of guava in surrounding regions in order to target high risk populations for immediate management consideration.

## Methods

### Fundamental and realized niches

Species distribution models for guava and guayabillo were constructed using MaxEnt v.3.4.1 (Phillips et al., 2006). For each species, a list of collections was compiled using herbarium collections from the Charles Darwin Foundation Herbarium (CDS), the National Herbarium of Ecuador (QCNE), and collections from previous work in this system (Reatini, 2021; Urquía et al., 2019, 2020). This included a total of 308 presence records for guayabillo and 336 for guava. Sampling bias was accounted for using the target background method following Phillips et al. (2009) using a weighted Gaussian distance kernel based on 924 presence records from all study systems within our collaborative dataset on interactions between invasive and endemic species pairs in the Galápagos. This dataset includes species from *Psidium, Lantana, Pennisetum, Cenchrus, Passiflora*, and *Gossypium*. Environmental data included the 19 bioclimatic variables from the WorldClim dataset (Hijmans et al., 2005), a digital elevation model (DEM) of the Galápagos (Souris, M. Institut de Recherche Pour Le Développement (IRD)), and a Galápagos ecosystems map (Galapagos Science Center General Resource). All layers were rescaled to a 50 meter resolution to match the spatial scale of the DEM for use with MaxEnt. Initial model performance was evaluated using the area under the receiver operating characteristic curve (AUC) for a test dataset consisting of a random subsample of 10% of the occurrence data for each species. The optimal regularization multiplier was set to 1 for both species after testing values of 1, 2, 5, and 10, with model selection performed using minimum Akaike Information Criterion (AIC) in ENMTools v1.3 (Warren et al., 2010) following Morales et al. (2017). Correlation filtering was then performed by removing one of each pair of variables with Pearson correlation coefficient >|0.7| to reduce model complexity and overfitting. The resulting six variables were the same for both species, and included four climate variables (Mean Diurnal Range, Annual Temperature Range, Annual Precipitation, and Precipitation of Coldest Quarter), the DEM, and the ecosystem layer. Final model performance was evaluated using AUC. Binary range maps of suitable habitat were generated using the 10th percentile training presence cloglog threshold for both species (57.9% probability of occurrence for guayabillo and 56.6% for guava) – which is a commonly used MaxEnt threshold method (Kramer-Schadt et al., 2013). Niche overlap ranging from 0 (completely different) to 1 (completely the same) was estimated using Warren’s *I* statistic in ENMTools v1.3 (Warren et al., 2010). Spatial overlap between binary range maps was used to classify predicted sympatry and allopatry for both species, and these were compared with the presence records above and with land cover maps from Laso et al. (2020) to compare our predictions of the fundamental niches (SDMs) and realized niches (land cover) of each species.

### Local exclusion on San Cristóbal Island

Due to the logistical difficulty of measuring survivorship for long-lived tree species, reproductive fitness alone was used to quantify population fitness on San Cristóbal Island. Fitness was estimated by measuring fruit set and seed set in sympatry and allopatry for both species. Specifically, fitness measurements were performed within two sympatric-allopatric site pairs (one pair on the east side of the island and one on the west side) for guayabillo, and one sympatric-allopatric site pair on the east side of the island for guava (Table S1).

For fruit set, branches with open flowers and mature flower buds were marked and the number of flowers and buds were recorded. After 20 days, the number of developing fruits on marked branches was recorded, and fruit set was measured as the proportion of flowers that produced fruit. Fruit set between sympatric and allopatric sites for both species was compared using generalized linear models with a logit link functions. Given that fruit set measurements were not available from allopatric CR due to permitting complications during the fruiting season, fruit set for guayabillo was compared between allopatric GAL and the two sympatric sites independently in addition to a comparison between sympatric and allopatric sites.

For seed set, 30 mature fruits were collected at random from each population and seed set was measured as the number of seeds within each fruit. Seed set for guayabillo was analyzed using an aligned ranks transformation ANOVA due to the non-normality of residuals and nonhomogeneity of variance, including as fixed factors patry (sympatry vs. allopatry), direction (eastern vs. western site pairs), and the interaction between patry and direction. For guava, seed set was analyzed using a one-way ANOVA due to normality of residuals and homogeneity of variance.

Tree height was used as a rough proxy for age to estimate the age structure of each population. For sites that contained guayabillo, three 10 m^2^ quadrats centered around randomly selected adult guayabillo individuals were analyzed. For allopatric EJ, quadrats were centered around randomly selected adult guava individuals. Within each quadrat, the number of individuals and the height of each individual were recorded for both species. In order to evaluate whether there was a deficit of guayabillo seedlings and juveniles in sympatry, the height distribution of guayabillo was then compared within each sympatric-allopatric site pair using Kolmogorov-Smirnov tests.

### Regional decline across the archipelago

To evaluate the extent of regional decline across the archipelago for guayabillo, the distribution of guayabillo populations throughout predicted sympatry was quantified for each island using ArcGIS Pro v.3.0.0. Specifically, predicted sympatry on each island was defined by extracting the overlap between the binary range maps for both species. The geographic centroid of predicted sympatry was then calculated for each island. A transect originating from the centroid of predicted sympatry was then drawn through each guayabillo population to the outer edge of predicted sympatry, and the total distance from the centroid to the edge and the distance between the population and the edge were recorded. The position of each of these populations was then normalized by dividing the distance between the point and edge by the total distance between the centroid and point, yielding the relative position between 0-1 of each population between the centroid (0) and edge (1) of predicted sympatry. Collections had to be separated by at least 200 meters to be considered a separate population for the sake of this analysis to account for potential sampling bias. To assess whether guayabillo has been significantly displaced from sympatry on San Cristóbal Island relative to islands with more recent introductions of guava, the distribution of guayabillo populations throughout predicted sympatry was compared between islands using a Kruskal-Wallis test followed by a post-hoc Conover-Iman test with Bonferroni correction for multiple comparisons using a family-wise error rate of 0.05 and per-test α of 0.05/6= 0.0083.

The extent of guava occupation in predicted sympatry was also quantified by calculating the percentage of overlap between predicted sympatry and the sum of guava dominant forests and mixed forests from available landcover maps (Laso et al., 2020). To account for the limited geographic extent of landcover maps, for this analysis we only considered the region of predicted sympatry that fell within the geographic extent of the available landcover map for each island.

### Risk assessment based on guava cover

Guava density was quantified in regions surrounding guayabillo populations from the regional decline dataset by cropping landcover maps from Laso et al. (2020) around those populations. Specifically, shape files of 1 km^2^ quadrats were centered around each population in ArcGIS Pro v.3.0.0 and landcover maps were trimmed around those quadrats. The percentage of land cover within each quadrat was then quantified including the cover of guava, mixed forest, other invasive species, cropland and pasture, native vegetation, and cover with no vegetation (*i*.*e*., built environment, bare ground, fresh water, *etc*.). Quadrats were sorted by guava density and plotted for each island, and these data were used to identify guayabillo populations at highest risk of exclusion for immediate management consideration.

## Results

### Fundamental and realized niches

Species distribution models (SDMs) were used to test whether guayabillo and guava’s predicted fundamental niches (the SDMs) and realized niches (contemporary distributions) exhibit patterns consistent with local exclusion of guayabillo by guava. The predicted fundamental niches of the two species were quite similar overall (*I*=0.868), and there was a large degree of overlap between the predicted ranges of both species on San Cristóbal Island – including much of the highlands (Figure 1A). Yet all extant guayabillo populations are limited to the outer edge of predicted sympatry or beyond (Figure 1A,E), which corresponds with the outer edge of guava’s realized range (Figure 1B). Thus, guava occupies the region where guayabillo is predicted to occur but does not.

### Local exclusion on San Cristóbal Island

To test whether sympatric guayabillo populations at lower elevations on San Cristóbal are in decline, fruit set, seed set, and seedling recruitment were all evaluated in sympatry relative to allopatry. Both fruit set and seed set were significantly lower in sympatry relative to allopatry for guayabillo (*P* = 0.001 and *P* = 2.78E-09 respectively; Table 1, Table S2). There was a significant difference in seed set between the eastern and western site pairs (*P* = 1.88E-06), and there was a significant interaction between patry and the site pairs (*P* = 0.001; Table S2) which appears to be driven by seed set being much lower in sympatric LG in the east relative to sympatric CM in the west (Figure 1C). The difference between the site pairs could not be evaluated for fruit set, where we have data for only one pair. For guava, fruit set was significantly lower in sympatric LG relative to allopatric EJ (*P* = 0.0197), but the opposite was true for seed set (*P* = 0.005; Table S3).

**Table 1.** Population level fruit set for guava and guayabillo in sympatry and allopatry.

| **Guayabillo** | **n** | **Fruit set (%)** |
| --- | --- | --- |
| LG (sympatric) | 205 | 12.2 |
| GAL (allopatric) | 156 | 20.51 |
| CM (sympatric) | 95 | 4.21 |
| CR (allopatric) | - | - |
| **Guava** |  |  |
| LG (sympatric) | 65 | 18.46 |
| EJ (allopatric) | 83 | 36.14 |

Using height as a rough proxy for age, there was a significant difference in the height distribution of guayabillo individuals between sympatry and allopatry for both site pairs (*P* =0.0169 in the west and *P*=0.0015 in the east; Table S4), which appears to be driven by a deficit of seedlings and juveniles in both sympatric sites (Figure 1D). In fact, only two guayabillo individuals <2 meters in height were found across both sympatric sites, whereas individuals <2m were abundant at both allopatric sites (38 and 34 individuals for GAL and CR respectively, Figure 1C). By contrast, guava individuals <2m were abundant in the sympatric site LG (57 individuals); of these, roughly half were confirmed to be asexually propagated (Figure S2B).However, only three guava individuals <2m were observed in the sympatric site CM (Figure S2A).

### Regional decline across the archipelago

To test whether the islands with a more recent guava invasion show a weaker sign of regional decline for guayabillo populations, the distribution of guayabillo populations throughout predicted sympatry was assessed for each coinhabited island. The distribution of guayabillo populations throughout predicted sympatry on San Cristóbal was significantly reduced relative to the three islands where guava was introduced more recently (Table S5, Figure 1E). On Santa Cruz island, where guava was introduced ca. 1950s (Urquía et al., 2019, 2021), guayabillo populations show the weakest signs of regional decline. Those on Floreana island, where guava was introduced ca. 1930s (Urquía et al., 2019, 2021), show intermediate signs of regional decline (Figure 1E) – although some degree of decline is apparent on all islands. The exact date of introduction on Isabela is unknown, but the available historic and genetic evidence suggests an intermediate introduction similar to Floreana (Urquía et al., 2019, 2021), which is consistent with the intermediate level of regional decline of guayabillo seen on that island (Figure 1E). However, the distribution of guayabillo throughout the area of predicted sympatry did not differ significantly between Floreana, Isabela, and Santa Cruz. Notably, for all islands except Isabela, the percentage of predicted sympatry occupied by guava corresponds roughly with the date of introduction – with earlier introductions having a greater extent of invasion (Figure S5). A massive wildfire on Isabela in 1994 decimated the highlands and was followed by rapid guava encroachment in the last two decades (Shimizu, 1997), which likely explains why Isabela has an unusually extensive invasion for its date of introduction (Figure S5).

### Risk assessment based on guava cover

Overall, guava density in the regions surrounding guayabillo populations varied considerably on each island (Figure 1F, Table 2). Populations with greater than 25% guava cover were present on all four islands, and populations with greater than 50% guava cover were found on San Cristóbal and Isabela. On San Cristóbal Island, guava density was highest in the region surrounding sympatric LG and decreased with elevation (Figure 1F).

**Table 2.** Guayabillo populations ranked by the sum of the percent cover of guava forest and mixed forest within quadrats for management consideration. LG (Rank 7) and CM (Rank 28) are bold to indicate risk categories.

| Rank | Population ID | latitude of center | longitude of center | Island | Guava | Mixed | Sum |
| --- | --- | --- | --- | --- | --- | --- | --- |
| 1 | I4 | -0.80548 | -91.0488 | Isabela | 50.25059 | 30.2305 | 80.48109 |
| 2 | I6 | -0.80348 | -91.0543 | Isabela | 44.09824 | 34.84303 | 78.94127 |
| 3 | I1 | -0.82355 | -91.0037 | Isabela | 75.08416 | 0.528075 | 75.61224 |
| 4 | I2 | -0.82248 | -91.0058 | Isabela | 67.69389 | 7.04786 | 74.74175 |
| 5 | I7 | -0.80479 | -91.053 | Isabela | 41.02546 | 32.91974 | 73.9452 |
| 6 | I3 | -0.8254 | -91.0024 | Isabela | 66.59943 | 0.750216 | 67.34964 |
| 7 | **LG** | **-0.8747** | **-89.4437** | **San Cristobal** | **54.50516** | **10.1177** | **64.62286** |
| 8 | I5 | -0.8082 | -91.0444 | Isabela | 44.50707 | 17.9073 | 62.41438 |
| 9 | I17 | -0.87458 | -91.0068 | Isabela | 0.418105 | 59.13827 | 59.55637 |
| 10 | F1 | -1.30175 | -90.4563 | Floreana | 37.89157 | 19.51069 | 57.40225 |
| 11 | I9 | -0.82408 | -91.019 | Isabela | 21.39422 | 26.86876 | 48.26299 |
| 12 | I13 | -0.87663 | -91.0082 | Isabela | 0.913618 | 44.66275 | 45.57637 |
| 13 | I22 | -0.85513 | -90.9929 | Isabela | 0.0378 | 40.72453 | 40.76233 |
| 14 | I20 | -0.87265 | -91.0023 | Isabela | 0.04337 | 40.46414 | 40.50751 |
| 15 | I8 | -0.81585 | -91.0355 | Isabela | 23.408 | 15.50185 | 38.90985 |
| 16 | SC1 | -0.88071 | -89.4364 | San Cristobal | 15.68346 | 22.26642 | 37.94987 |
| 17 | I21 | -0.8533 | -90.9913 | Isabela | 0.0378 | 36.31772 | 36.35552 |
| 18 | SZ40 | -0.7029 | -90.3433 | Santa Cruz | 0.048097 | 35.83804 | 35.88614 |
| 19 | I10 | -0.83517 | -91.001 | Isabela | 8.084115 | 27.25156 | 35.33567 |
| 20 | SC6 | -0.90695 | -89.4419 | San Cristobal | 0.848306 | 34.16887 | 35.01717 |
| 21 | I11 | -0.83605 | -91.0005 | Isabela | 6.2661 | 28.58247 | 34.84857 |
| 22 | F2 | -1.31294 | -90.4484 | Floreana | 14.3008 | 19.05276 | 33.35355 |
| 23 | SZ1 | -0.63888 | -90.2899 | Santa Cruz | 33.04294 | 0.1953 | 33.23824 |
| 24 | SC7 | -0.90957 | -89.4412 | San Cristobal | 0.335079 | 30.10627 | 30.44134 |
| 25 | I18 | -0.85543 | -91 | Isabela | 0.216 | 30.1324 | 30.3484 |
| 26 | SZ2 | -0.64272 | -90.2847 | Santa Cruz | 29.10715 | 0.603983 | 29.71114 |
| 27 | SZ3 | -0.6429 | -90.2881 | Santa Cruz | 29.06216 | 0.1953 | 29.25746 |
| 28 | **CM** | **-0.90495** | **-89.5682** | **San Cristobal** | **2.74223** | **26.3406** | **29.08288** |
| 29 | SZ4 | -0.67512 | -90.3398 | Santa Cruz | 26.84701 | 0.437479 | 27.28449 |
| 30 | SC2 | -0.89295 | -89.4374 | San Cristobal | 6.919103 | 20.01387 | 26.93297 |
| 31 | SZ5 | -0.63652 | -90.2913 | Santa Cruz | 25.70149 | 0.171 | 25.87249 |
| 32 | F3 | -1.31492 | -90.4328 | Floreana | 5.386004 | 19.11869 | 24.50469 |
| 33 | SC5 | -0.9231 | -89.473 | San Cristobal | 2.069595 | 22.00986 | 24.07946 |
| 34 | SC4 | -0.92359 | -89.4664 | San Cristobal | 3.433098 | 20.0118 | 23.4449 |
| 35 | I15 | -0.85352 | -91.0009 | Isabela | 0.63818 | 21.70845 | 22.34663 |
| 36 | SC3 | -0.89094 | -89.4377 | San Cristobal | 4.997275 | 16.89626 | 21.89353 |
| 37 | I16 | -0.85512 | -91.002 | Isabela | 0.495 | 20.90942 | 21.40442 |
| 38 | SZ6 | -0.67718 | -90.34 | Santa Cruz | 20.48721 | 0.7326 | 21.21981 |
| 39 | SZ7 | -0.65768 | -90.4089 | Santa Cruz | 19.753 | 1.390631 | 21.14363 |
| 40 | SZ8 | -0.63878 | -90.2938 | Santa Cruz | 18.62619 | 2.514681 | 21.14088 |
| 41 | SZ16 | -0.69858 | -90.4033 | Santa Cruz | 9.800983 | 11.20897 | 21.00995 |
| 42 | SZ15 | -0.69633 | -90.4008 | Santa Cruz | 10.84971 | 9.496335 | 20.34605 |
| 43 | SZ28 | -0.67245 | -90.2706 | Santa Cruz | 2.730972 | 16.9353 | 19.66627 |
| 44 | SZ29 | -0.69012 | -90.3162 | Santa Cruz | 2.696182 | 15.94132 | 18.63751 |
| 45 | SZ26 | -0.67027 | -90.2702 | Santa Cruz | 3.386835 | 14.76538 | 18.15222 |
| 46 | SZ10 | -0.67227 | -90.438 | Santa Cruz | 14.66281 | 1.975189 | 16.638 |
| 47 | I12 | -0.82094 | -91.026 | Isabela | 4.50443 | 12.08944 | 16.59387 |
| 48 | F4 | -1.31614 | -90.4526 | Floreana | 3.95968 | 12.44714 | 16.40682 |
| 49 | SZ9 | -0.67032 | -90.4354 | Santa Cruz | 14.8572 | 0.6345 | 15.4917 |
| 50 | SZ12 | -0.68712 | -90.4135 | Santa Cruz | 12.59515 | 2.620167 | 15.21532 |
| 51 | SZ13 | -0.63005 | -90.4366 | Santa Cruz | 12.04367 | 3.170856 | 15.21453 |
| 52 | I19 | -0.84207 | -90.9913 | Isabela | 0.213716 | 13.67237 | 13.88609 |
| 53 | SZ11 | -0.65071 | -90.2891 | Santa Cruz | 12.5959 | 1.230806 | 13.82671 |
| 54 | I14 | -0.81868 | -91.0226 | Isabela | 0.821104 | 11.77278 | 12.59388 |
| 55 | SZ17 | -0.66567 | -90.2754 | Santa Cruz | 8.345628 | 3.79062 | 12.13625 |
| 56 | SZ14 | -0.63287 | -90.4358 | Santa Cruz | 10.9846 | 0.325617 | 11.31022 |
| 57 | SZ36 | -0.70795 | -90.3374 | Santa Cruz | 0.911816 | 10.2128 | 11.12461 |
| 58 | SZ20 | -0.70808 | -90.3561 | Santa Cruz | 6.232704 | 3.565856 | 9.79856 |
| 59 | SZ21 | -0.70785 | -90.3519 | Santa Cruz | 5.978137 | 3.76354 | 9.741677 |
| 60 | SZ23 | -0.70768 | -90.3488 | Santa Cruz | 4.475929 | 4.57376 | 9.049688 |
| 61 | SZ18 | -0.66863 | -90.439 | Santa Cruz | 7.594206 | 0.651165 | 8.245371 |
| 62 | SZ19 | -0.66632 | -90.4394 | Santa Cruz | 7.331486 | 0.4302 | 7.761686 |
| 63 | SZ22 | -0.7029 | -90.3433 | Santa Cruz | 4.747925 | 2.787922 | 7.535847 |
| 64 | SZ38 | -0.70783 | -90.3164 | Santa Cruz | 0.435786 | 6.242736 | 6.678522 |
| 65 | SZ31 | -0.67512 | -90.3398 | Santa Cruz | 1.943877 | 4.432319 | 6.376196 |
| 66 | SZ24 | -0.70088 | -90.3416 | Santa Cruz | 3.804296 | 2.539117 | 6.343414 |
| 67 | SZ27 | -0.70477 | -90.3437 | Santa Cruz | 3.070105 | 3.072739 | 6.142844 |
| 68 | SZ25 | -0.69818 | -90.3406 | Santa Cruz | 3.576031 | 1.380781 | 4.956812 |
| 69 | SZ33 | -0.70657 | -90.3217 | Santa Cruz | 1.429952 | 3.221868 | 4.65182 |
| 70 | SZ30 | -0.69012 | -90.3162 | Santa Cruz | 2.683335 | 1.259376 | 3.942711 |
| 71 | SZ37 | -0.69273 | -90.3136 | Santa Cruz | 0.51067 | 2.991644 | 3.502314 |
| 72 | SZ32 | -0.6939 | -90.3213 | Santa Cruz | 1.735594 | 1.265434 | 3.001028 |
| 73 | SZ35 | -0.69263 | -90.3173 | Santa Cruz | 1.207244 | 1.3698 | 2.577044 |
| 74 | SZ34 | -0.6948 | -90.3194 | Santa Cruz | 1.413807 | 0.644791 | 2.058599 |
| 75 | SZ39 | -0.70932 | -90.3203 | Santa Cruz | 0.2826 | 0.762537 | 1.045137 |

## Discussion

Using a combination of species distribution modeling, population fitness metrics, and geospatial analyses, we tested the hypothesis that invasive guava has excluded endemic guayabillo from the highlands of San Cristóbal Island, and that sympatric populations at lower elevations – and on islands with more recent guava introductions – may currently be under threat of exclusion.

Our species distribution analyses reveal patterns consistent with historical exclusion of guayabillo on San Cristóbal by guava. Specifically, extant guayabillo populations are limited to the outer edge of predicted sympatry on San Cristóbal Island, and guava dominated forests occupy much of predicted sympatry where guayabillo is now absent (Figure 1A,B). Therefore, guayabillo’s realized niche is a subset of its fundamental niche, and interactions with guava appear to explain the difference. These results paint a clear picture of recent replacement of guayabillo populations by invasive guava.

Our analyses of reproductive fitness and seedling recruitment in sympatry corroborate the strong association of guava with the decline of guayabillo on San Cristóbal. Specifically, the significant reduction in both fruit set and seed set in sympatric guayabillo populations relative to allopatric populations indicates that the presence of invasive guava is associated with low reproductive output of guayabillo populations (Figure 1C, Table 1, Table S2). Reduced reproductive fitness can lead to demographic decline and ultimately to local extirpation if the number of offspring produced fall below the number required for replacement. This appears to be the case, given the deficit of seedlings and juveniles in sympatry relative to allopatry and consequently the significant difference in the height distribution between sites (Figure 1D, Table S4). Together, these results suggest that the presence of guava is either a direct causal agent of the ongoing demographic decline of sympatric guayabillo populations, or that both the expansion of guava and decline of guayabillo are driven by a common factor or factors.

Our geospatial analyses of regional decline offer an additional strong line of evidence linking the spread of guava to the decline of guayabillo. The distribution of guayabillo populations throughout predicted sympatry on San Cristóbal Island is significantly reduced relative to islands with more recent introductions of guava. Moreover, there is a striking relationship between the date of guava’s introduction and the extent of guayabillo’s regional decline across the four coinhabited islands (Figure 1E). Although the extent of decline is most advanced on San Cristóbal, some degree of regional decline is evident on all four coinhabited islands (Figure 1E).

### What biotic interactions could be driving guayabillo’s decline?

Although the results presented here cannot identify the precise mechanism(s) of guayabillo’s decline, they are consistent with the idea that direct competition or negative sexual interactions (reproductive interference) between guava and guayabillo may be among the contributing factors. First, guava is known to negatively impact a broad range of native species in other regions (including in island ecosystems) via resource competition (CABI, 2022), and our estimations of the fundamental niches of guava and guayabillo show a large degree of niche overlap (Warren’s *I*=0.868), which is a favorable condition for resource competition. Second, reproductive interference is predicted to be relatively common during secondary contact between native and introduced species (Kyogoku, 2020), and this system features characteristics that would make reproductive interference likely. Specifically, the prevalence of generalist pollination networks in the Galápagos promotes heterospecific mating between native and introduced species (Olesen et al., 2002), there is a general lack of intrinsic prezygotic barriers observed between guava and other *Psidium* species (Cardoso et al., 2017; Gomes et al., 2017; Landrum et al., 1995), and there is a difference in ploidy between diploid guava and polyploid guayabillo (Reatini, 2021) which can generate strong postzygotic barriers – as observed between guava and other polyploid *Psidium* species (Cardoso et al., 2017; Gomes et al., 2017). Further work will be required to more directly test the role of resource competition, reproductive interference, and other possible causal factors in driving the decline of guayabillo.

### Conservation implications and recommendations for management

Our results suggest that sympatric guayabillo populations across the archipelago are likely to suffer the same fate as those on San Cristóbal without management intervention, but also provide a few critical pieces of information to help guide management decisions and prevent further decline of this protected Galápagos endemic. If biotic interactions with guava are indeed the causal factor driving guayabillo’s decline, then this has a number of implications. First, control of guava could help the guayabillo populations to recover. Second, the density of guava surrounding a guayabillo population can be used to indicate of the level of risk for local extirpation of guayabillo, and this can be measured from remote sensing data. This would provide a powerful tool for land managers by offering a means of prioritizing management targets. Given that LG shows strong evidence of imminent exclusion (i.e. drastically reduced reproductive fitness and seedling recruitment; Figure 1C, D, Table 1, Table S2), we recommend that guayabillo populations in regions with similar or greater guava cover than LG (sum of guava forest and mixed forest cover >64.6%) should be the highest priority targets for immediate conservation action. Seven such populations exist, LG on San Cristóbal, and 6 populations in the highlands of Isabela (Table 2) – which notably are located where the wildfire encouraged rapid encroachment by guava. Given that we still see evidence for demographic decline (albeit weaker) in CM, we use CM as the lower bound for moderate risk of imminent exclusion, and we recommend management of the resulting 20 moderate risk populations in ranked order following Table 2. We classify the remaining populations with lower guava cover than CM as the lowest risk category. However, it is important to note that these populations are still at risk of exclusion given that they are still sympatric populations within the range of habitat classified as suitable for guava in our SDMs, and risk may increase if guava density rises in these regions. Thus, the results we present here not only reveal an ongoing conservation threat, but also provide a tool to guide immediate conservation action in order to prevent further decline of this vulnerable Galápagos endemic tree species.

## Acknowledgements

We thank the Galápagos National Park, the Galápagos Science Center, the Center for Galápagos Studies, and the Charles Darwin Foundation for providing the logistical and administrative support that made this work possible. BR has been supported by the Fulbright U.S. Student Program, an R. C. Lewontin award from the Society for the Study of Evolution, and the UNC Chapel Hill Department of Biology. This work was also supported by a grant from the UNC Center for Galápagos Studies to TJV.

## Supporting Information

**Figure S1.**
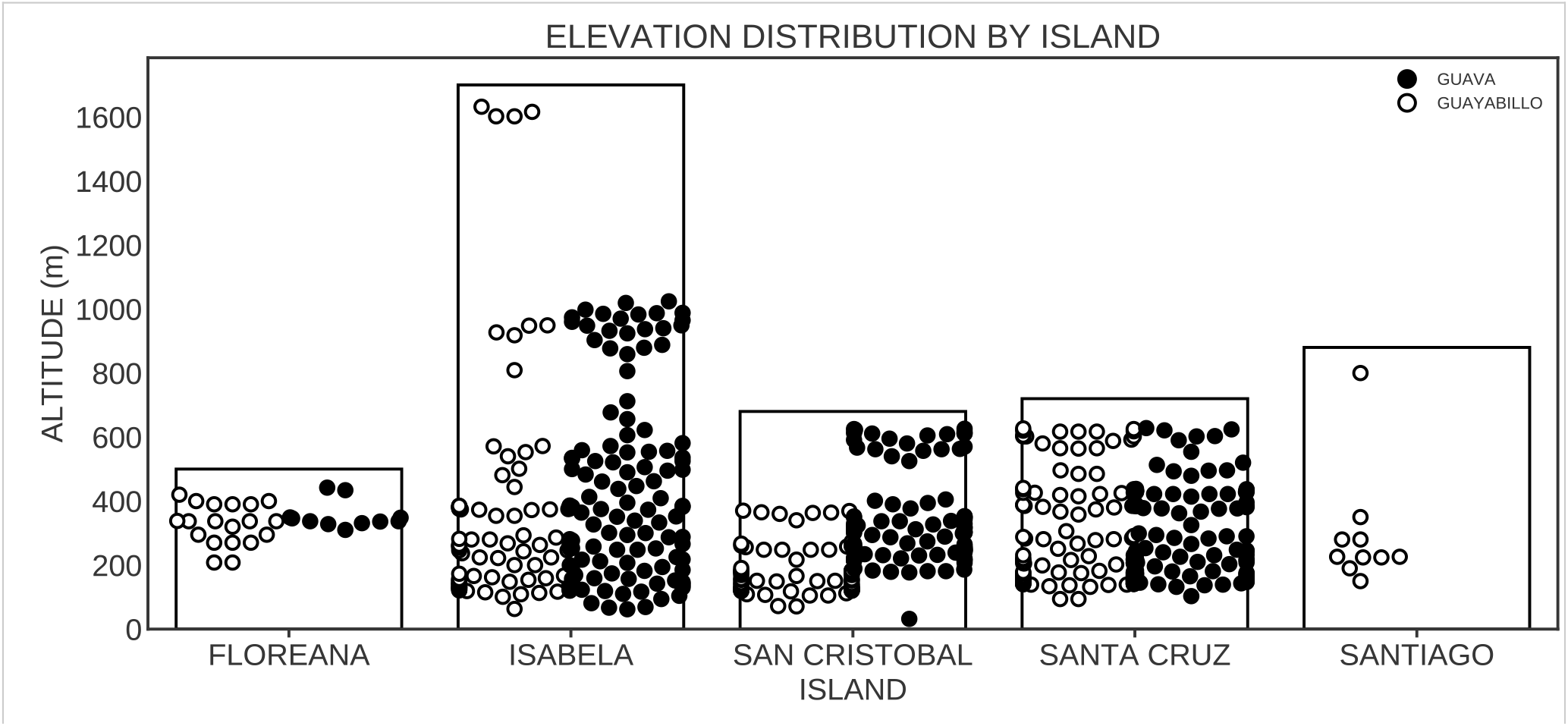
Guava and guayabillo elevation distribution. Individuals of guayabillo (open circles) are found in the highlands of all islands except San Cristóbal, whereas those of guava (filled circles) are found in the highlands of all four coinhabited islands.

**Figure S2.**
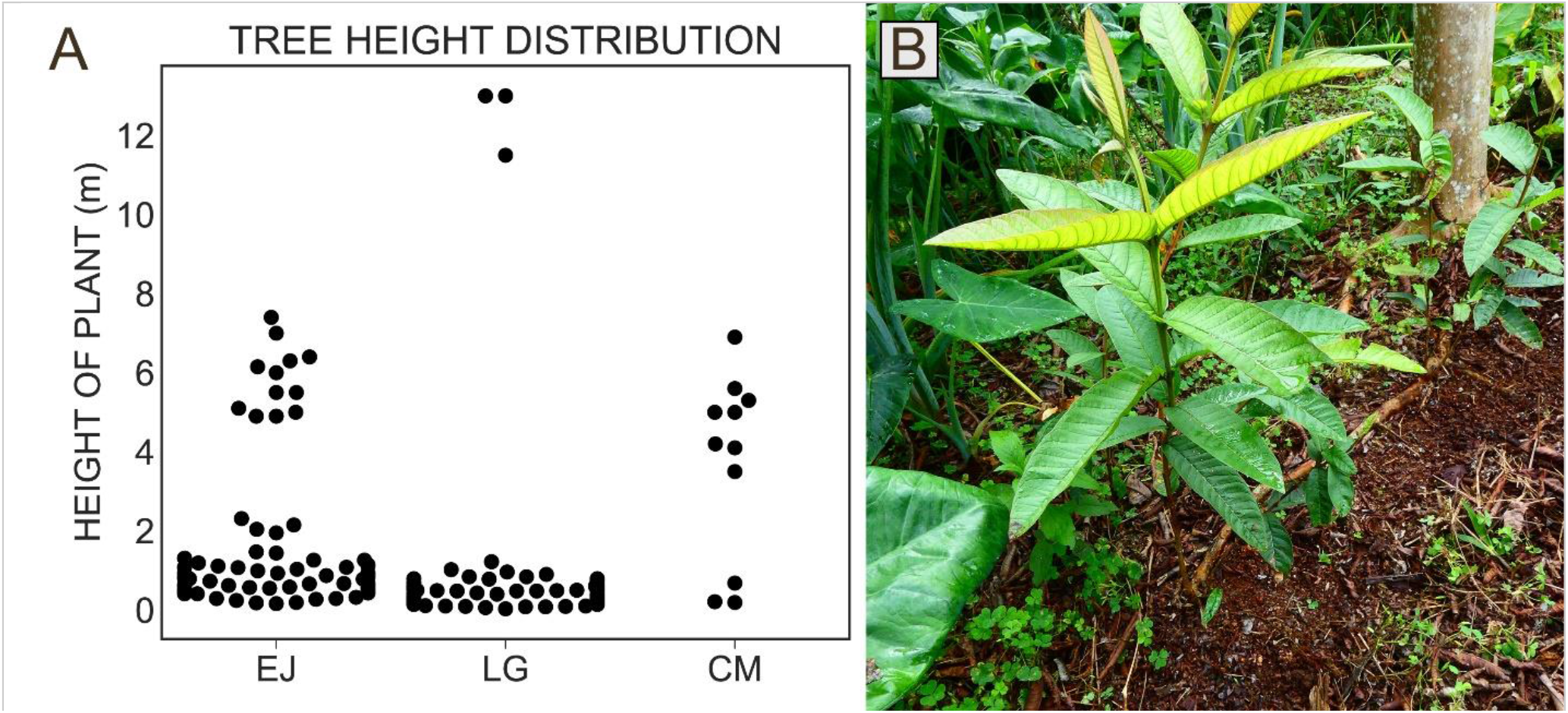
Age proxy data and image of clonal reproduction. A) Height distribution as a proxy for age of guava individuals in allopatric EJ and sympatric LG and CM shows that fewer juveniles are present in CM, perhaps because CM is located at the invasion front in an area of mixed forest where guava is not yet the most predominant species. B) Clonal reproduction of guava observed in sympatric LG suggests that the viability of a guava population does not depend entirely upon sexual reproduction.

**Figure S3.**
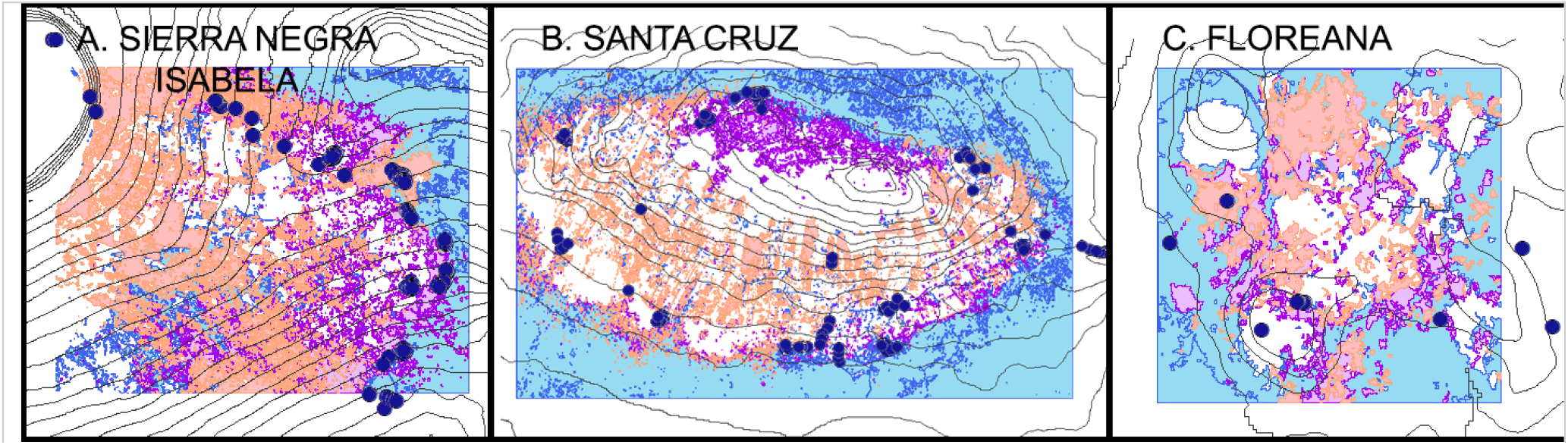
Land cover maps from Laso et al. (2020) for the coinhabited islands of Isabela, Santa Cruz, and Floreana. Extant guayabillo populations (large dots) located in guava dominated forests (pink) on all coinhabited islands are at high risk of extirpation. Guayabillo populations within mixed forests where guava is present but not yet the predominant species (purple), and within native forests (blue) are currently at lower risk of extirpation by guava.

**Figure S4.**
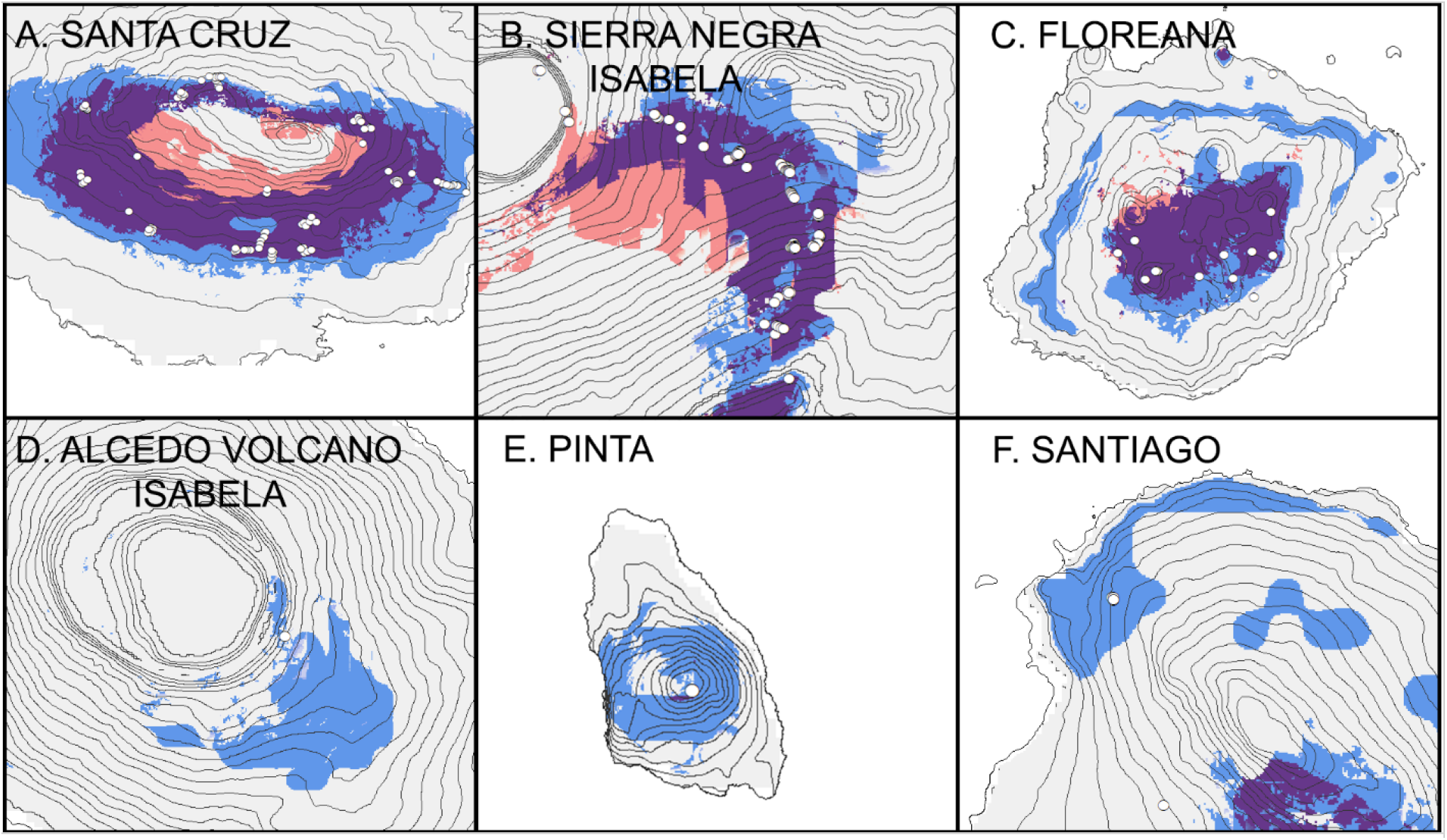
Predicted ranges for guava (pink) and guayabillo (blue) including predicted sympatry (purple). Isolated guayabillo populations (white circles) are rarely present in sympatry except at the periphery of guava’s predicted range. (A-D) coinhabited islands, (E and F) isolated islands.

**Figure S5.**
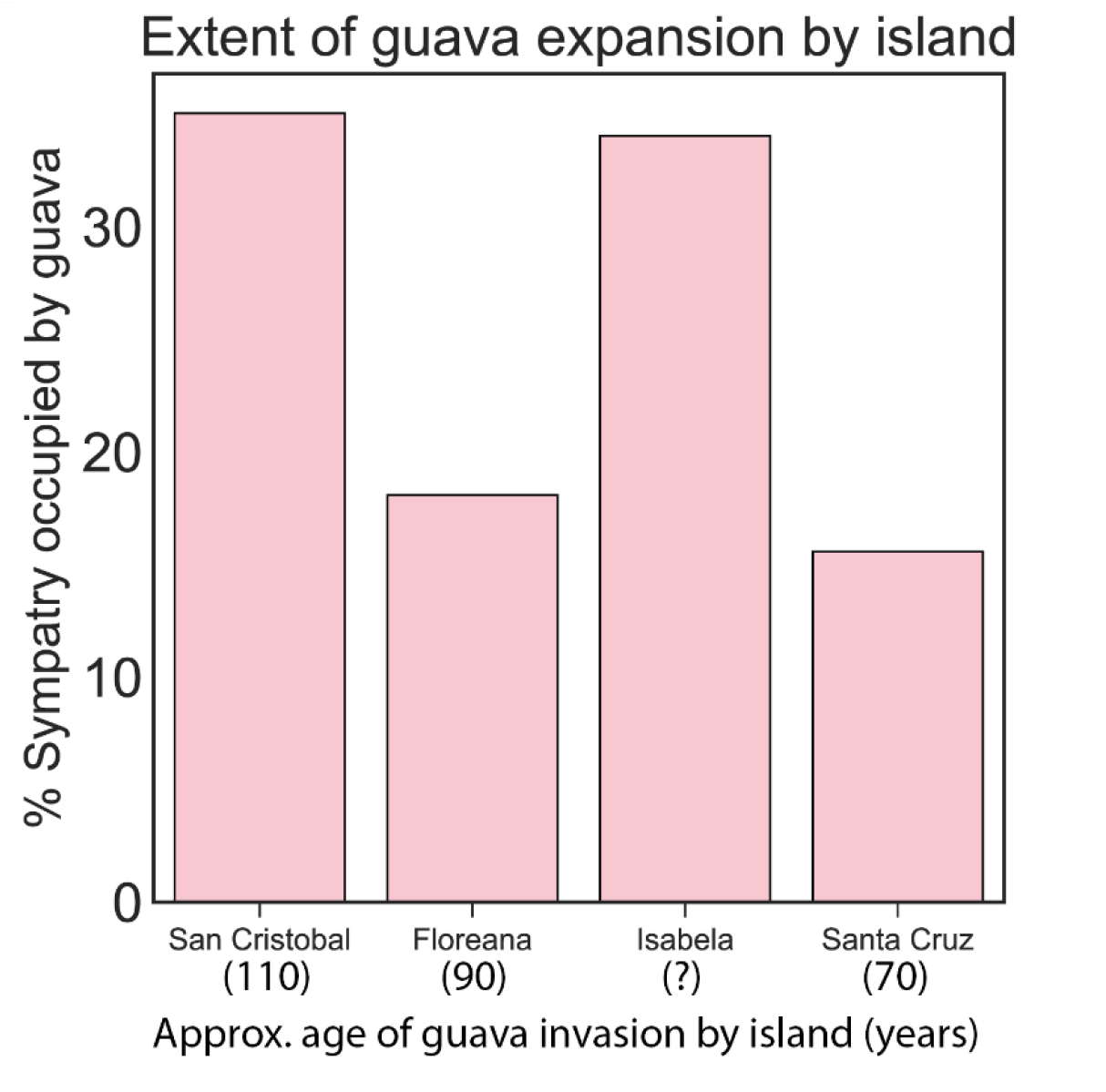
Percentage of predicted sympatry occupied by the guava invasion on each coinhabited island. With the exception of Isabela Island, the extent of the guava invasion is greater on islands with earlier introductions than on islands with more recent introductions.

**Table S1.** Study sites on San Cristóbal.

| Site name | species present | Altitude<br>(m) | Latitude<br>(degrees S) | Longitude<br>(degrees W) |
| --- | --- | --- | --- | --- |
| El Junco (EJ) | *guava* | 600 | -0.89055 | -89.48144 |
| Las Goteras (LG) | both | 360 | -0.87449 | -89.44407 |
| Cerro Mundo (CM) | both | 248 | -0.90203 | -89.56916 |
| Centro de Reciclaje<br>(CR) | *guayabillo* | 178 | -0.91306 | -89.57863 |
| Galapaguera (GAL) | *guayabillo* | 158 | -0.91364 | -89.43716 |

**Table S2.** The effect of sympatry vs. allopatry on reproductive success in guayabillo.

|  |  |  |  |  |
| --- | --- | --- | --- | --- |
| Guayabillo Seed Set<sup>1</sup> | degrees of freedom (df) | df.residuals | *F* | *P* |
| patry | 1 | 116 | 41.504 | 2.78E-09 |
| direction | 1 | 116 | 25.214 | 1.88E-06 |
| patry:direction | 1 | 116 | 10.985 | 0.001226 |
| Guayabillo Fruit Set<sup>2</sup> | estimate | standard error | *z* | *P* |
| (Intercept) | -1.3545 | 0.1983 | -6.832 | 8.40E-12 |
| patry | -0.8876 | 0.2783 | -3.189 | 0.00143 |
| GAL:CM | -1.77 | 0.5479 | -3.23 | 0.00124 |
| GAL:LG | -0.6306 | 0.2912 | -2.165 | 0.03036 |
| <sup>1</sup>Analysis of Variance of aligned rank transformed data. |  |  |  |  |
| <sup>2</sup>Generalized Linear Model with logit link function. |  |  |  |  |

**Table S3.**
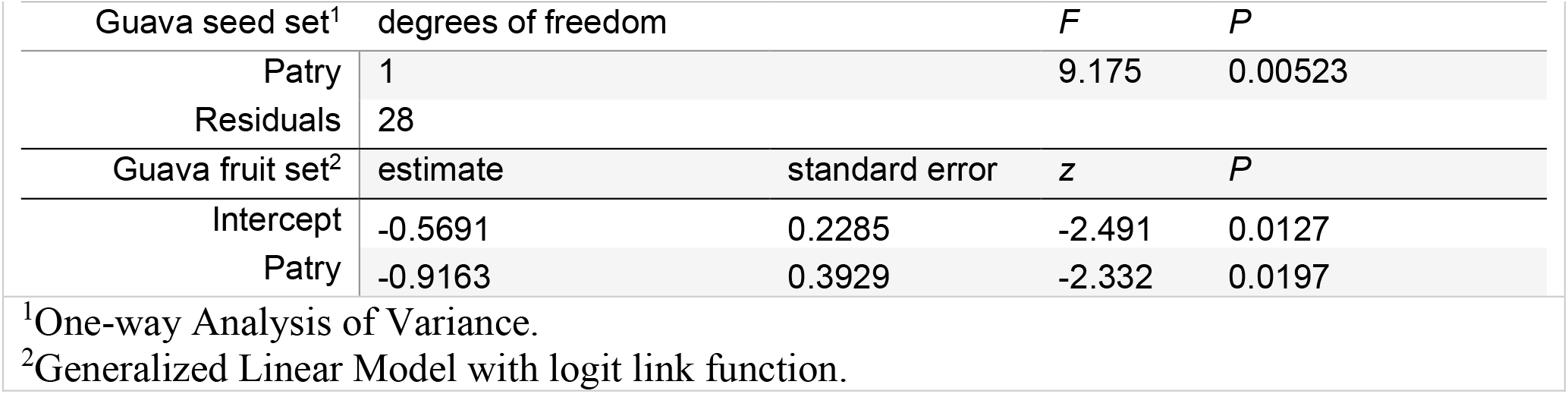
The effect of sympatry vs. allopatry on reproductive success in guava.

**Table S4.** The effect of sympatry vs. allopatry on height distribution in guayabillo.

|  |  |  |
| --- | --- | --- |
| Guayabillo height<sup>1</sup> | *D* | *P* |
| CM-CR | 0.49371 | 0.01696 |
| LG-GAL | 0.98077 | 0.001577 |
| <sup>1</sup>Two-way Kolmogorov-Smirnov tests |  |  |

**Table S5.** The distribution of guayabillo populations within predicted sympatry.

|  |  |  |  |
| --- | --- | --- | --- |
| Kruskal-Wallis test | $\chi^2$ | *df* | *P* |
| Archipelago-wide | 15.816 | 3 | 0.001237 |
| *Post hoc* Conover-Iman tests | *t* |  | *P* |
| Floreana - Isabela | 0.656158 |  | 0.2568 |
| Floreana - San Cristobal | 2.940600 |  | 0.0021* |
| Isabela - San Cristobal | 2.862220 |  | 0.0027* |
| Floreana - Santa Cruz | -0.605743 |  | 0.2732 |
| Isabela - Santa Cruz | -1.826064 |  | 0.0357 |
| San Cristobal - Santa Cruz | -4.250829 |  | 0.0001* |
| *Asterisks mark significant comparisons following Bonferroni correction. |  |  |  |

